# Force and stepwise movements of gliding motility in human pathogenic bacterium *Mycoplasma pneumoniae*

**DOI:** 10.1101/2021.05.25.445585

**Authors:** Masaki Mizutani, Yuya Sasajima, Makoto Miyata

**Affiliations:** Graduate School of Science, Osaka City University, 3-3-138 Sugimoto, Sumiyoshi-ku, Osaka 558-8585, Japan; The OCU Advanced Research Institute for Natural Science and Technology (OCARINA), Osaka City University, 3-3-138 Sugimoto, Sumiyoshi-ku, Osaka 558-8585, Japan

**Keywords:** Motility, Optical tweezers, class *Mollicutes*, Infection

## Abstract

*Mycoplasma pneumoniae*, a human pathogenic bacterium, binds to sialylated oligosaccharides and glides on host cell surfaces via a unique mechanism. Gliding motility is essential for initiating the infectious process. In the present study, we measured the stall force of an *M. pneumoniae* cell carrying a bead that was manipulated using optical tweezers on two strains. The stall forces of M129 and FH strains were averaged to be 23.7 and 19.7 pN, respectively, much weaker than those of other bacterial surface motilities. The binding activity and gliding speed of the M129 strain on sialylated oligosaccharides were eight and two times higher than those of the FH strain, respectively, showing that binding activity is not linked to gliding force. Gliding speed decreased when cell binding was reduced by addition of free sialylated oligosaccharides, indicating the existence of a drag force during gliding. We detected stepwise movements, likely caused by a single leg under 0.2–0.3 mM free sialylated oligosaccharides. A step size of 14–19 nm showed that 25–35 propulsion steps per second are required to achieve the usual gliding speed. The step size was reduced to less than half with the load applied using optical tweezers, showing that a 2.5 pN force from a cell is exerted on a leg. The work performed in this step was 16%–30% of the free energy of the hydrolysis of ATP molecules, suggesting that this step is linked to the elementary process of *M. pneumoniae* gliding.

**IMPORTANCE:** Human mycoplasma pneumonia is caused by the bacterium *Mycoplasma pneumoniae*. This tiny bacterium, shaped like a missile, binds to human epithelial surfaces and spreads using a unique gliding mechanism to establish infection. Here, we analyzed the movements and force of this motility using a special setup: optical tweezers. We then obtained detailed mechanical data to understand this mechanism. Furthermore, we succeeded in detecting small steps of nanometers in its gliding, which is likely linked to the elementary process of the core reaction: chemical to mechanical energy conversion. These data provide critical information to both control this human pathogen and explore new ideas for artificial molecular machines.

## INTRODUCTION

Members of the bacterial class *Mollicutes*, which includes the genus *Mycoplasma*, are parasitic and occasionally commensal bacteria that are characterized by small cells and genomes and by the absence of a peptidoglycan layer (1, 2). *Mycoplasma* species bind to host cell surfaces and exhibit gliding motility to spread the infectious area. Interestingly, *Mycoplasma* gliding does not involve flagella or pili and is completely unrelated to other bacterial motility systems or conventional motor proteins that are common in eukaryotic motility (3). The gliding motility of *Mycoplasma* is divided into two types, *Mycoplasma pneumoniae* and *Mycoplasma mobile*, which share no homology in component proteins, indicating independent mechanisms (4, 5). *M. pneumoniae* is a human pathogen that causes respiratory diseases, including bronchitis and atypical pneumonia (6). *M. pneumoniae* cells bind to human epithelial surfaces through sialylated oligosaccharides (SOs), which are major structures of animal cell surfaces related to cell-cell recognition, and the binding targets of many pathogens and toxins (7–10). The cells show unidirectional gliding motility at a speed of up to 1 μm/s on SO-coated glass surfaces (Fig. 1A) (5, 11), which is known to be essential for their infection (12). The gliding machinery, called the “attachment organelle,” is localized at a cell pole (13). The attachment organelle is divided into two parts: internal and surface structures. The internal structure is composed of an internal core complex and a surrounding translucent area. The internal core comprises three parts: a terminal button, paired plates, and a bowl complex from the front side of the cell (Fig. 1B) (5, 14, 15). The major surface structure, called “P1 adhesin complex” or “genitalium and pneumoniae cytoadhesin (GPCA)”, is composed of P1 adhesin and P40/P90 proteins and aligned around the internal structure, which plays a dual role as the adhesin to bind to SOs and as the leg for gliding (Fig. 1B) (16–20). The model for gliding called the “Inchworm model” or “Double-spring hybrid ratchet model” is proposed, in which cells repeat the extensions and contractions of the attachment organelle based on the energy from ATP hydrolysis to enable smooth gliding (15, 21–23). Generally, the mechanical characteristics and detailed analysis of movements are essential for creating and completing a detailed model for the motility mechanism (24–28). However, to date, no information is available about the force for gliding.

**FIG 1.**
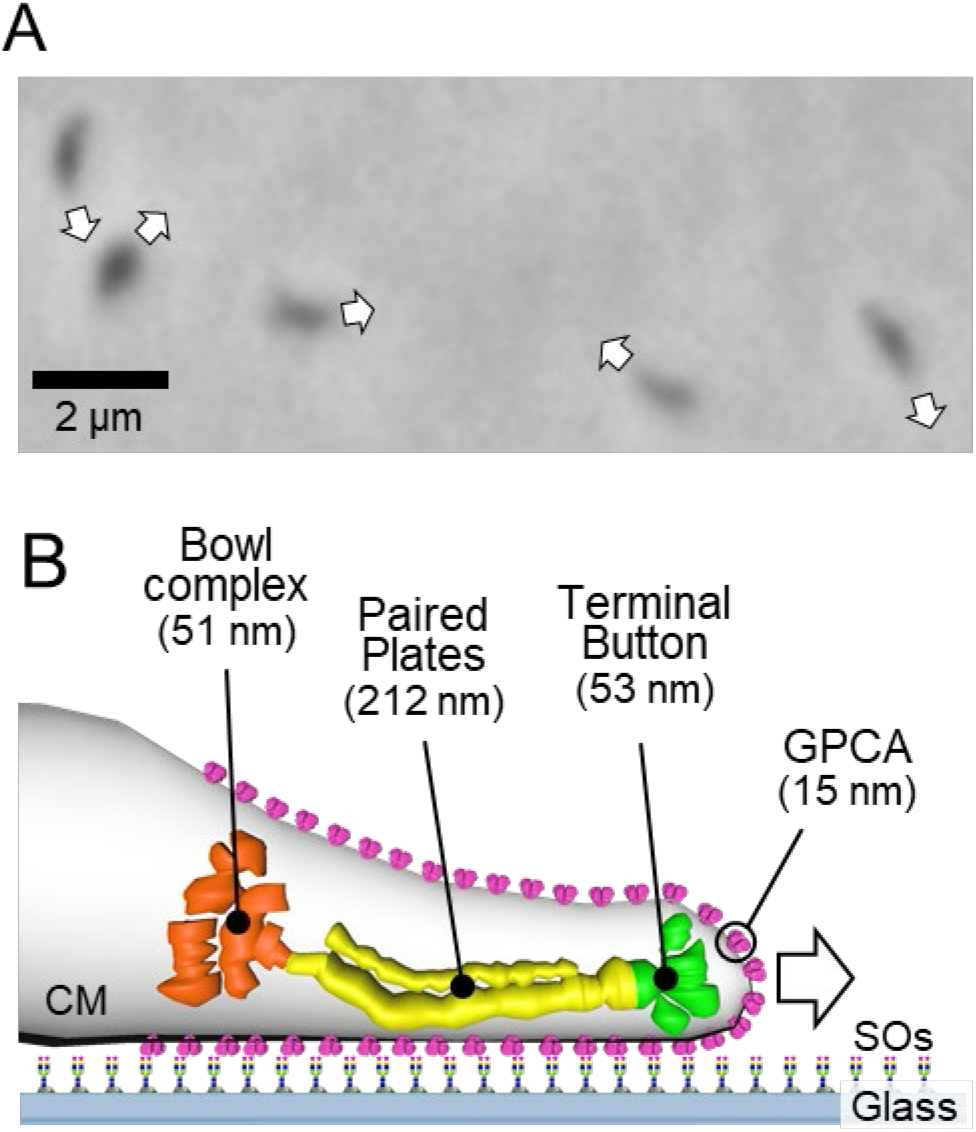
Gliding motility of *Mycoplasma pneumoniae*. (A) Phase-contrast micrograph of *M. pneumoniae* cells. The gliding directions of cells are indicated by white arrows. (B) Illustration of gliding machinery. The gliding machinery is composed of the bowl complex, paired plates, terminal button, and GPCA. The lengths along the cell axis are summarized from previous studies (14, 19). GPCA, working as a leg, is anchored to the cell membrane (CM) and catches sialylated oligosaccharides (SOs) fixed on solid surfaces.

In the present study, we measured the stall forces of the two strains and discuss the relationship between binding and force. Furthermore, we succeeded in detecting and measuring stepwise movements that are likely linked to the elementary process of the gliding reaction.

## Results

### Stall force measurement using optical tweezers

Optical tweezers are commonly used to measure the stall force generated by pili in bacterial motility or motor proteins in eukaryotic motility (29–32). The stall force is defined as the force needed to stop movements and is equal to the maximal propulsion force for locomotion. Previously, we measured the stall force for the gliding motility of *M. mobile*, based on a mechanism unrelated to *M. pneumoniae* gliding, using optical tweezers (33, 34). In the present study, we applied this method to *M. pneumoniae* gliding. *M. pneumoniae* cells were biotinylated and inserted into a tunnel chamber, which was assembled using two glass plates and double-sided tape (35). An avidin-conjugated polystyrene bead was trapped by a highly focused laser beam and attached to a gliding cell at the back end of the cell body by exploiting the avidin-biotin interaction. The cells pulled the bead from the trap center with gliding and then stalled (Fig. 2A and B; *SI Appendix*, Movie S1). The force was calculated by measuring the distance between the centers of the bead and the laser trap, which was multiplied by the trap stiffness; the force acting on the bead increased linearly with the displacement from the trap center (29). Starting from 0 s, the pulling force increased and reached a plateau in 120 s (Fig. 2C). The maximal value of the force averaged over one second was determined as the stall force. The stall force of M129 strain cells was 23.7 ± 6.3 pN (Fig. 2E). We also measured the stall force of the FH strain, another major strain of *M. pneumoniae*. The cells of the FH strain pulled the bead in a manner similar to that of M129 cells (*SI Appendix*, Movie S2). The stall force of the FH strain was 19.7 ± 5.3 pN, significantly weaker than that of the M129 strain (Fig. 2D and E).

**FIG 2.**
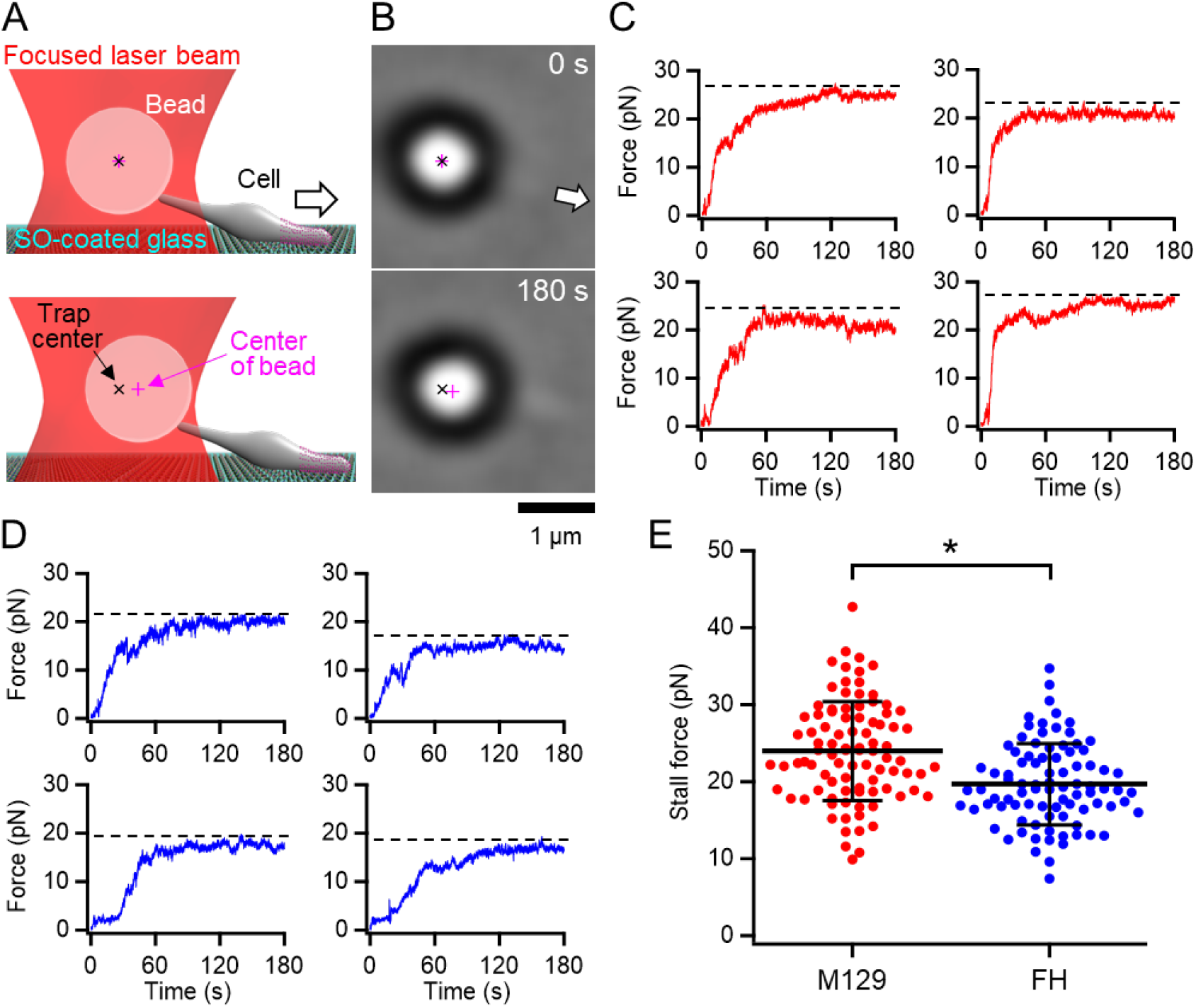
Stall force measurements. (A) Illustrations of experimental design for force measurements. The white arrow indicates the gliding direction. (B) Optical micrographs of a trapped cell. The cell attached with a bead (large black ring with white center) glided in the direction of the white arrow from 0 and stalled in 180 s. The cell image can be observed near the base of the arrow. (C) Four representative time courses of force increments for M129 strain. The broken lines indicate the values of stall force. (D) Four representative time courses of force increments for FH strain. The broken lines indicate the values of stall force. (E) Scatter dot plot of stall force (*n* = 92 and 88 in M129 and FH, respectively) shown with averages (thick lines) and standard deviations (thin lines). \**p* = 2.4×10^−6^ by Student’s *t*-test.

### Binding activity and gliding motility

To characterize the differences in gliding motility, we measured the binding activity and gliding speed of the M129 and FH strains. *M. pneumoniae* cells were suspended in HEPES buffer containing 20 mM glucose to obtain an optical density at 595 nm of 0.07, then inserted into tunnel chambers. After incubation for 5–30 min, the tunnel chambers were washed and observed by phase-contrast microscopy. The number of bound cells in the M129 strain increased with time from 90 ± 16 to 308 ± 20 cells in 100 ×100 μm^2^ area from 5 to 30 min (Fig. 3A and B). These values are consistent with the results of a previous report (9). In contrast, the number of bound cells in the FH strain increased from 12 ± 2 to 36 ± 8 cells from 5 to 30 min (Fig. 3A and B). The number of bound cells in the FH strain was 7.2–8.7-fold smaller than that of the M129 strain at all time points (Fig. 3B). When 10× concentrated cell suspension was examined, the bound cell numbers of the FH strain increased from 117 ± 10 to 386 ± 24 cells from 5 to 30 min, 1.2—1.3 times that of M129 at all time points (Fig. 3A and B). These results indicate that the FH strain has approximately 8-fold lower binding activity to SO-coated glass surfaces than the M129 strain. Considering that the stall force of FH was only 1.2-fold smaller than that of the M129 strain, the force is unlikely to be linked to binding activity.

**FIG 3.**
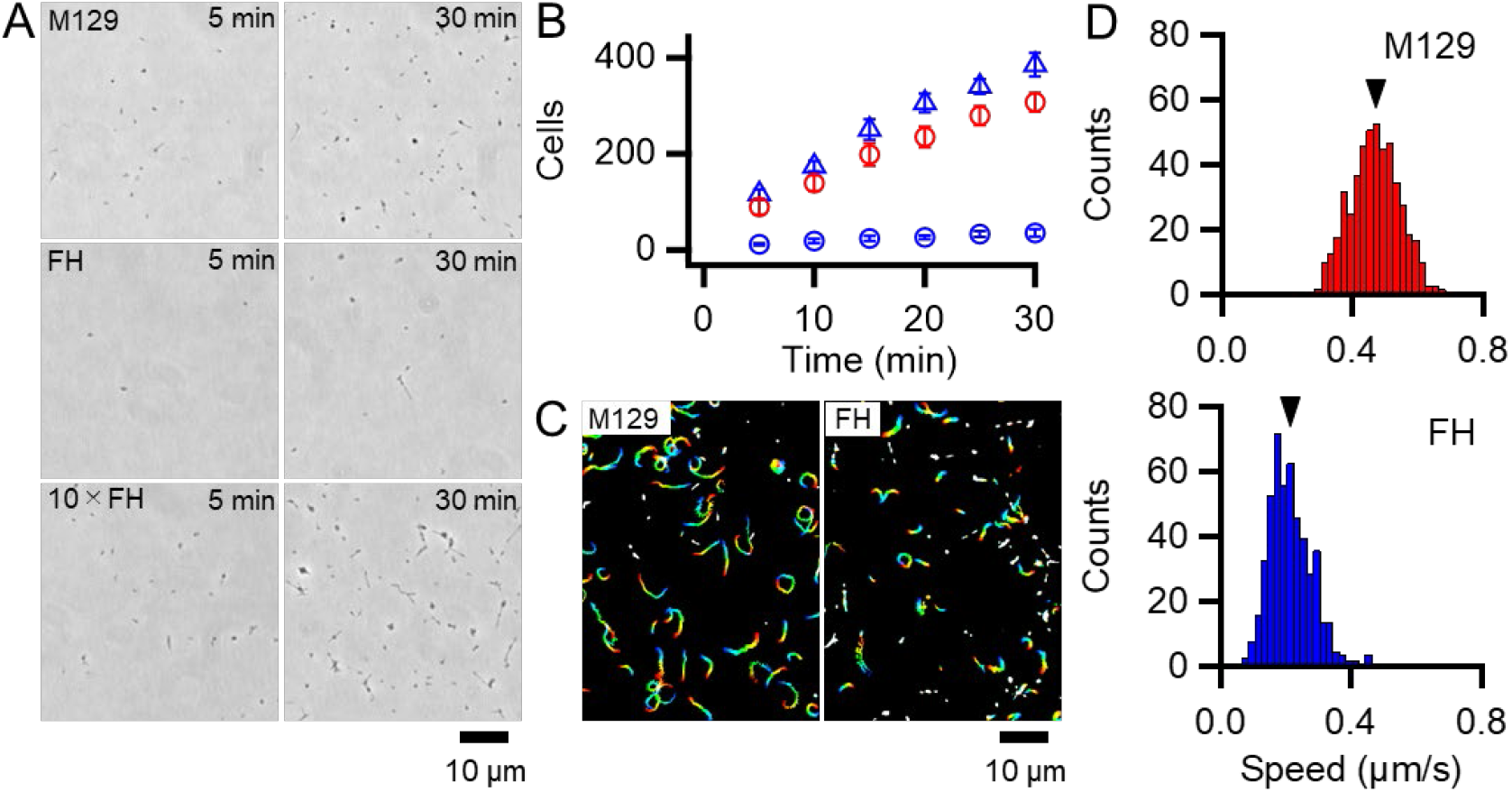
Binding activity and gliding motility. (A) Phase-contrast micrographs of cells on SO-coated glass surfaces. M129, FH, and 10× concentrated FH are shown after 5 and 30 min of incubation. (B) Averaged bound cell numbers in 100×100 μm^2^ plotted with standard deviations (*n* = 10 in each). M129 (red open circle), FH (blue open circle), and 10× concentrated FH (blue open triangle) are presented. (C) Cell trajectories for 20 s, changing color with time from red to blue. (D) Distributions of gliding speeds averaged for 20 s at 1-s intervals (*n* = 500 in each). Mean values are marked by black triangles.

Most of the bound cells showed gliding motility on the SO-coated glass (Fig. 3C; *SI Appendix*, Movies S3 and S4). To characterize gliding motility, the proportion of gliding cells and gliding speed were analyzed. The proportion of gliding cells to all bound cells was 78.1% (*n_total_* = 2284, *n_glide_* = 1785) and 60.0% (*n_total_* = 2271, *n_glide_* = 1362) in the M129 and FH strains, respectively (Fig. 3C). The gliding speeds averaged for 20 s at 1-s intervals were 0.47 ± 0.08 and 0.21 ± 0.07 μm/s for M129 and FH strains, respectively (Fig. 3D). These results suggest that binding activity is not directly linked to gliding speed.

### Differences in amino acid sequences of gliding proteins between two strains

The FH strain used in this study has not been genomically analyzed. Therefore, we sequenced the genomes of both strains using MiSeq (*SI Appendix*, Dataset S1) and analyzed the sequences of 14 genes that have been reported to be involved in binding and gliding (5). Only one amino acid was substituted in the M129 strain genome from a standard genome, M129-B7 (GenBank accession no. CP003913), which was V196A in the HMW3 protein, which is positioned at the terminal button in the attachment organelle (Fig. 1B). Our FH strain had 161 and 155 variations from the reported FH and FH2009 strains, respectively (GenBank accession no. CP010546 and CP017327), indicating that the FH strain is distant from the genome strains. The differences between the two strains analyzed in the present study in terms of the 14 genes were as follows: (i) P1 adhesin showed many differences, including 87 single amino acid substitutions, 18 amino acid insertions, and four amino acid deletions, known differences of FH from M129 strains (*SI Appendix*, Fig. S1). (ii) The P40/P90 protein showed 96 single amino acid substitutions, one amino acid insertion, and 68 amino acid deletions (*SI Appendix*, Fig. S2). (iii) The other 12 genes showed 0–5 mutations in each gene, as summarized in *SI Appendix*, Table S1.

### Effect of free sialyllactose on binding and gliding

*M. pneumoniae* cells have hundreds of legs and glide smoothly on solid surfaces. To trace the behavior of a single propulsion event, the number of working legs should be reduced by adding a free form of SO, which is the binding target for gliding (9, 34, 36). To quantitatively analyze the effects of free SO on the binding activity and gliding under our conditions, we measured the bound ratio of cells and the gliding speed of the M129 strain under various concentrations of the SO, 3’-*N*-acetylneuraminyllactose (SL). *M. pneumoniae* cells suspended in the buffer were inserted into a tunnel chamber. Then, the cell behavior in 0–0.5 mM SL solutions was analyzed. The addition of free SL slowed down and then stopped gliding or released the gliding cells from the glass surfaces, but did not release non-gliding cells, indicating that release requires glass binding by GPCA with displacements (Fig. 4A). The number of gliding cells and the gliding speed relative to the initial speed were decreased by 0.1–0.5 mM with SL treatments from 55 ± 11% to 26 ± 16% and from 0.33 ± 0.06 to 0.19 ± 0.07 μm/s, respectively (Fig. 4B and C). We then decided to use 0.2–0.3 mM concentrations for further experiments because the binding and gliding were partially inhibited under these conditions, which were advantageous for observing single propulsion events caused by a single leg during gliding.

**FIG 4.**
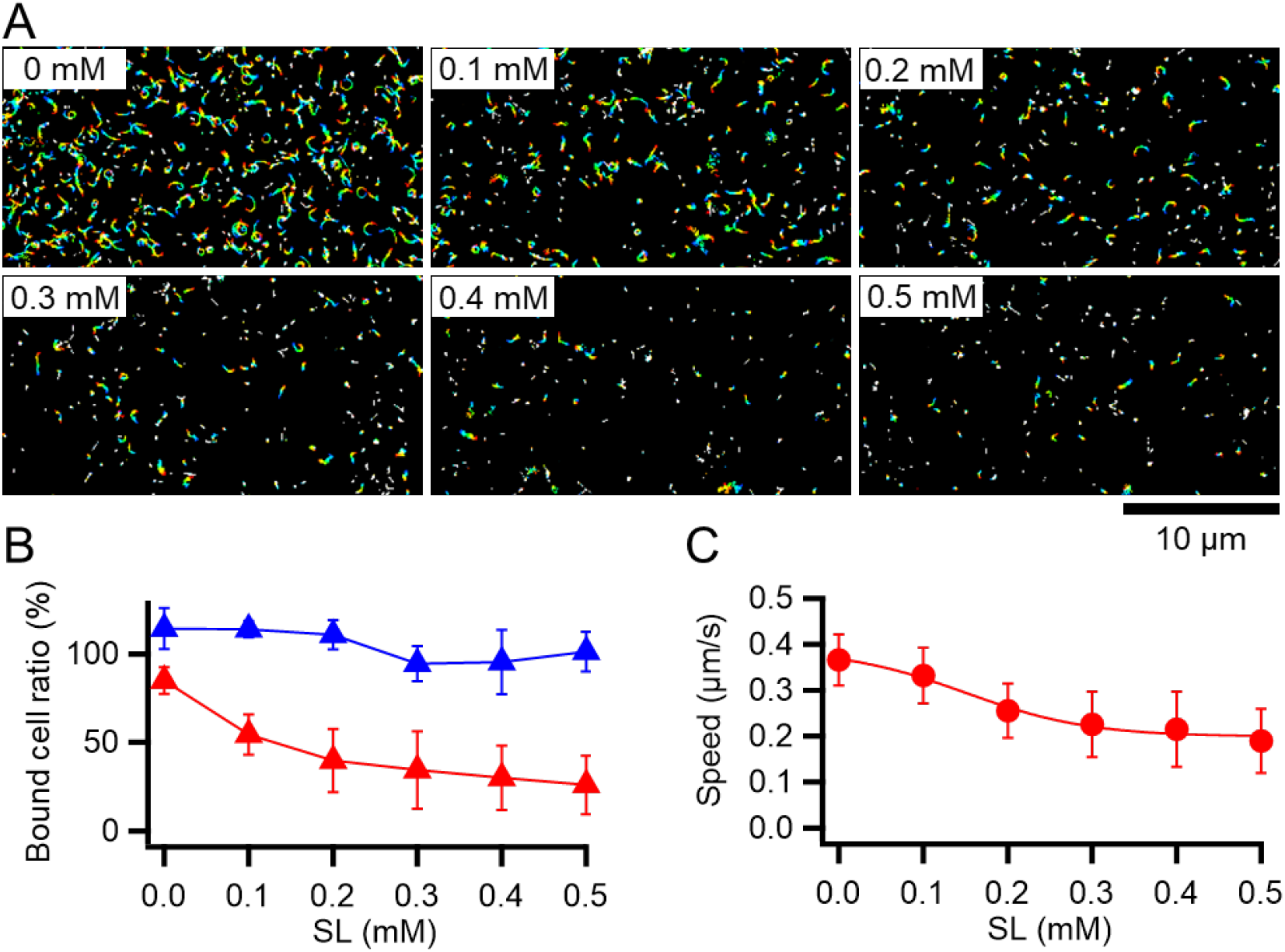
Effects of SL on binding and gliding. (A) Gliding cells were treated with free SL. The cell trajectories after SL treatment are presented as a stack for 20 s, changing color from red to blue. SL concentrations are shown at the left upper corner of each panel. (B) Number of cells bound to the glass at 5 min after treatment are plotted relative to the initial cell number. The averages for gliding (red triangles) and non-gliding (blue triangles) cells are shown with the standard deviations (*n* = 10, 10, 10, 10, 9, and 10 for 0, 0.1, 0.2, 0.3, 0.4, and 0.5 mM, respectively). (C) Averaged gliding speeds under various SL concentrations are plotted with the standard deviations and fitted by a sigmoidal curve (*n* = 50, 50, 50, 50, 56, and 56 for 0, 0.1, 0.2, 0.3, 0.4, and 0.5 mM, respectively).

### Stepwise gliding movements observed under free SL

Next, we traced cells under 0.2–0.3 mM free SL conditions as performed in the stall force measurements. The gliding cells slowly pulled the beads from the trap center. Half of the tested cells stalled in 120 s, but the other cells repeated creeping movements and detachment from the glass surfaces (Fig. 5A). The stall force of the cells that reached a plateau in 120 s was 16.2 ± 4.5 pN (Fig. 5B), which was significantly weaker than that without SL (*p* = 1.1×10^−5^ by Student’s *t*-test). In creeping movements, cells occasionally showed discontinuous displacements, which were mostly stepwise (Fig. 5C and D). Individual displacements in a stepwise time course were analyzed using the pairwise distance function (34, 36). The displacements shown in Fig. 5D were uniformly distributed at about 17-, 14-, and 16-nm intervals under 0.15, 0.20, and 0.17 pN/nm of trap stiffness, corresponding to 2.55, 2.80, and 2.66 pN of propulsion force, respectively (Fig. 5D and E). Note that force increments are generally calculated from the trap stiffness × displacement (29, 34).

**FIG 5.**
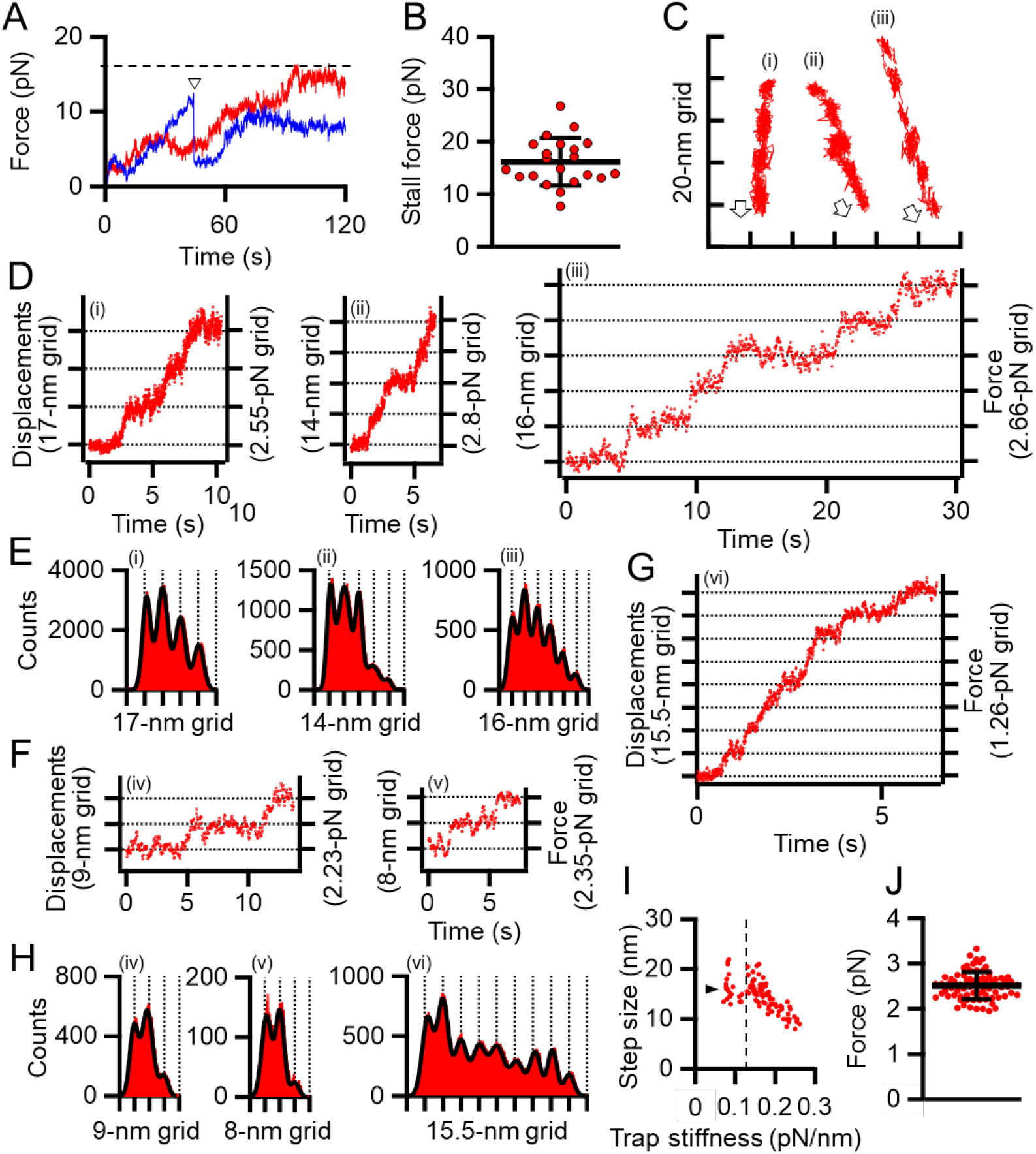
Stepwise movements under SL. (A) Representative force traces with time under 0.2–0.3 mM SL are shown for stalled (red) and detached (blue) cells. The broken line indicates the stall force. The time point of detachment is marked by a triangle. (B) Scatter dot plot of stall force shown with averages (thick line) and standard deviations (thin lines) (*n* = 21). (C) Three cell trajectories with stepwise movements are shown in a field. Open arrows indicate the gliding direction. (D) Displacement and force of cells whose trajectories are shown in panel (C). (E) Histograms of pairwise distance function (PDF) analysis of panel (D) were fitted by the sum of Gaussian curves. (F) Displacement and force under high trap stiffness. (G) Displacement and force under low trap stiffness. (H) Histograms of PDF analysis of panel (F) and (G). (I) Distribution of step size under various trap stiffnesses. Step sizes were plotted as a function of trap stiffness (*n* = 97). The position at 0.12 pN/nm of trap stiffness was marked by a dotted line. The average of step size under 0.07–0.12 pN/nm of trap stiffness was marked by a triangle. (J) Scatter dot plot of force increments under 0.13–0.26 pN/nm of trap stiffness shown with averages (thick line) and standard deviations (thin lines) (*n* = 76).

Next, to examine the load dependency of step sizes in detail, we measured step sizes under 0.07–0.26 pN/nm of trap stiffness. Ninety-seven steps in 36 cell trajectories, including at least two continuous steps, were detected. The step sizes under 0.13 to 0.26 pN/nm of trap stiffness linearly decreased from 21.0 to 8.0 nm with trap stiffness (Fig. 5F and H). In contrast, the step sizes from 0.07 to 0.12 pN/nm of trap stiffness were mostly constant and distributed at 14–19 nm with an average of 16.2 ± 2.7 nm (*n* = 21) (Fig. 5G–I).

From 0.07 to 0.12 pN/nm of trap stiffness, the force increments increased with the trap stiffness (Fig. 5I). In the cases where the force values increase with trap stiffness, the calculated force does not reflect the actual force because the load is too small to influence the movements. Therefore, we focused on the force increments measured under 0.13–0.26 pN/nm of trap stiffness. The force exerted in a single propulsion step was concluded to be 2.5 ± 0.3 pN (*n* = 76) (Fig. 5J).

## Discussion

### *M. pneumoniae* has weak stall force

In the present study, we measured the stall force in *M. pneumoniae* gliding using optical tweezers. Previously, the stall force was measured for some bacterial surface motilities. *Neisseria gonorrhoeae* and *Myxococcus xanthus* show surface motility driven by the retraction of type IV pili (37). The stall forces of single-pilus retraction in *N. gonorrhoeae* and *M. xanthus* were measured by optical tweezers to be approximately 80 and 150 pN, respectively (30, 38). *M. mobile* glides up to 4.0 μm/s by a mechanism unrelated to that of *M. pneumoniae*. Previously, we measured the stall force of *M. mobile* gliding using optical tweezers to be approximately 113 pN (23). The stall forces of *M. pneumoniae* gliding were 23.7 ± 6.3 and 19.7 ± 5.3 pN in M129 and FH strains, respectively (Fig. 2E), much weaker than those of other bacterial surface motilities. *M. pneumoniae* has a streamlined cell body about 0.2 μm in diameter (39). This shape and cell size may be beneficial for gliding in human tissues with weak forces.

In liquid culture, *M. pneumoniae* cells bind to the bottom surface of the tissue culture flask. This is distinct from the case of *M. mobile*, in which most cells float in the medium. Therefore, *M. pneumoniae* is expected to have a stronger force for gliding than *M. mobile*. However, we found that *M. pneumoniae* had a much weaker stall force than *M. mobile* (Fig. 2E). When comparing the two strains of *M. pneumoniae*, the FH strain showed much less active binding than, but a stall force comparable to, that of M129 (Fig. 2E and 3B). These facts indicate that the binding activity of *M. pneumoniae* cells is not determined by the gliding force, which is represented by the stall force.

### Drag force in gliding

The gliding speed decreased when cell binding was partially inhibited by the addition of free SL (Fig. 4). This observation is consistent with previous data; that is, the inhibition of binding by monoclonal antibodies decreased the gliding speed of *M. pneumoniae* (40) and the inhibition by SL decreased the speed of *M. gallisepticum*, coinciding with the common mechanism with *M. pneumoniae* (23). The decrease in speed was probably caused by the drag force generated from the substrate surface, because the friction force exerted from water is estimated to be more than 5000 times smaller than the stall force of 24 pN (Fig. 2E) (41, 42). As the cause of the drag force, two possibilities are considered: GPCA and others. If some proportion of GPCA molecules are not involved in gliding, they are not released from SOs by inhibitory factors, resulting in speed reduction. Interestingly, these observations and explanations are similar to the case of *M. mobile* gliding, even though they do not share the same structure of machinery (4, 9, 42–44). This scheme may be advantageous for gliding on SOs based on ATP energy.

### Stepwise movement as elementary process

We succeeded in detecting the stepwise movements of *M. pneumoniae* gliding. *M. pneumoniae* cells glided at a speed of 0.47 μm/s (Fig. 3D), and the step size in the load-free condition was 14–19 nm (Fig. 5I), suggesting 25–35 steps per second. Stepwise movements are well-studied in ATP-driven eukaryotic motor proteins including myosin, dynein, and kinesin (26, 45). The force and displacement of a single step in myosin II, cytoplasmic dynein, kinesin-1, and myosin V have been reported as 3–5, 7–8, 8, and 2–3 pN and 5.3, 8, 8, and 36 nm, respectively (24, 29, 31, 32, 46, 47). Stepwise movements are also present in bacterial motility. A flagellar motor shows 14 degrees of revolution as a step (27, 48). The gliding of *M. mobile* shows stepwise movements of 1.5 pN force and 70 nm length (34, 36). Generally, stepwise movements are thought to correspond to the elementary process of a motility event.

Previously, we showed that *M. pneumoniae*-type gliding motility is driven by energy from ATP hydrolysis (23). Motilities driven by ATP energy can be divided into elementary processes that are directly coupled with ATP hydrolysis, such as stepwise movements. The elementary processes observed as steps require a smaller amount of work than the energy produced by ATP hydrolysis, which is ∼80 pN nm (49). Therefore, we estimated the work done in the stepwise movements of *M. pneumoniae* gliding to determine this possibility. The work was estimated to be 18.2 ± 5.4 pN nm from the equation *W*_step_ = 0.5 × spring constant × displacement^2^, under 0.13–0.26 pN/nm of trap stiffness, where the stiffness is large enough to determine the force (*SI Appendix*, Fig. S3) (34). This value is 16%–30% of the free energy of the hydrolysis of ATP molecules, suggesting that we detected the elementary process of *M. pneumoniae* gliding. The energy conversion efficiencies of stepwise movements are 12%–40%, 10%, and 40%–60% for myosin II, cytoplasmic dynein, and kinesin, respectively (26, 45). Previously, we estimated it to be 10%–40% for *M. mobile* gliding (34). These facts suggest that the energy efficiency of stepwise movements of *M. pneumoniae* is in a similar range of myosin II and *M. mobile* gliding.

### Load-dependent step size

*M. pneumoniae* cells showed different step sizes depending on the load provided by optical tweezers continuously (Fig. 5I), suggesting that the step size of *M. pneumoniae* is load-dependent. In the infectious process, *M. pneumoniae* cells glide to the deep positions of respiratory systems and experience a large load at these positions (12). The load-dependent stepping behavior would be useful to glide against large loads because a load-independent stepping motor, kinesin, shows frequent back steps under large loads (50, 51). Different step sizes under different loads have also been observed in *M. mobile* gliding (34, 36).

### Suggestion for gliding mechanism

How can we image the gliding mechanism? The GPCA probably plays a critical role in *M. pneumoniae* gliding, because antibodies against the P1 adhesin, a component of GPCA, decreased the gliding speed and ultimately replaced the *M. pneumoniae* cells on the glass surface (40). Recently, the detailed structure of GPCA has been solved (17–20). It is a mushroom structure composed of two P1 adhesins and two P40/P90 molecules. The C-terminal regions of the four molecules are bundled and anchored to the cell membrane. P40 and P90 proteins are synthesized as a single protein and processed into two proteins that are not observed in *M. genitalium*. Although P1 adhesin is believed to be the receptor for SOs, the binding site exists at the distal end of P40/P90. This complex is thought to undergo conformational changes between open and closed with respect to the binding pockets, which are likely involved in the gliding mechanism (18, 20).

The attachment organelle responsible for gliding can be divided into a surface structure including GPCA and an internal rod structure (15, 22, 52–54). Briefly, the force for gliding is likely generated at the bowl because a few proteins essential for gliding and not binding are localized there (55–57). The paired plates and elastic components play a role in force transmission, because the gliding speed decreases severely in a deletion mutant (58). The terminal button likely connects the rod front to the cell membrane, because an end component, the P30 protein, features transmembrane segments (59, 60).

Here, we focus on the following observations to construct the working model for the gliding scheme: (1) The force is probably generated around the bowl complex, transmits through the internal structures including paired plates, and reaches GPCAs. (2) Inhibition of cell binding decreases gliding speed. (3) GPCA has open and closed conformations. (4) Gliding can be divided into steps because binding and force are not tightly coupled. (5) The gliding movement can be divided into 14–19 nm steps with a 2.5 pN force.

### Gliding scheme to explain stepwise movements

The model is composed of repeated cycle of four stages (i)–(iv) connected by four steps: “Power stroke,” “Release,” “Displacement,” and “Catch” (Fig. 6). (i) The GPCA in the open state catches an SO on the glass surface. (ii) Contraction of the internal structure pulls the cell body with 2.5 pN and a step size of 14–19 nm. This step is possibly linked to energy from an ATP molecule. (iii) GPCA switches into a closed state by triggering the pulling force transmitted from other working GPCAs, resulting in the release of after-stroke GPCA from the SO. (iv) The released GPCA returns to the open state and displaces another SO in the next position by the extension of the internal structure. (i) The GPCA captures the next SO. The full cycle was then repeated. This scheme explains the relationship between the binding and force of different strains, if we assume that release is enhanced and power stroke and catch are reduced in FH compared with those in M129. The observation that shorter steps occur under load can be explained if we consider that the power stroke is shortened by the load (Fig. 4I). Previously, our group suggested a working model for *M. pneumoniae* gliding, focusing mainly on the information of the internal structure with regard to the attachment organelle (5, 15). The previous model suggested “directed detachment of feet” (GPCA), due to the lack of information about step, force, and foot structure. In this study, we succeeded in adding new information and completing an updated model.

**FIG 6.**
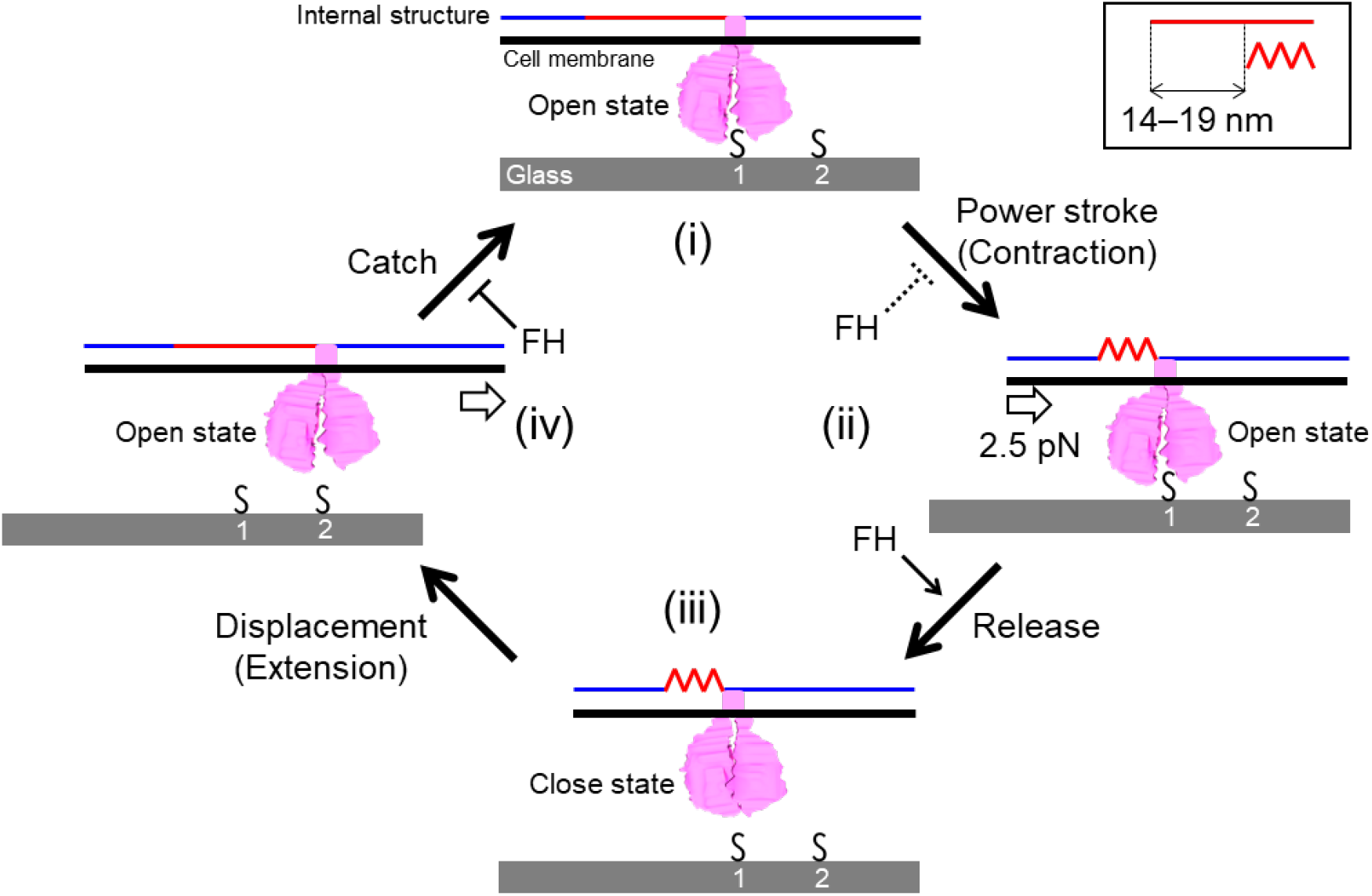
Schematic illustration of leg behaviors in stepwise movements. Internal structure, represented by blue and red lines, is responsible for force transmission. It repeats contraction and extension cooperatively with the surface structures. Conformational change of internal structure and step size are shown in the right-upper box. GPCA is presented in pink. Sialylated oligosaccharides are shown by “S.” Force exerted in a step is shown by an arrow in panel (ii). Displacement of the cell body forward is shown by an arrow in panel (iv). The gliding occurs through stages i to iv. See text for details.

## Materials and Methods

### Strains and Cultivation

*M. pneumoniae* M129 (ATCC29342) and FH strains were grown in SP-4 medium at 37°C in tissue culture flasks (TPP Techno Plastic Products AG, Trasadingen, Switzerland), as described previously (11, 61). The FH strain was kindly provided by Tsuyoshi Kenri at the National Institute of Infectious Diseases, Tokyo, Japan.

### Optical tweezers system

An inverted microscope (IX71; Olympus, Tokyo, Japan) was equipped with a Nd:YAG laser (ASF1JE01; Furukawa Electric, Tokyo, Japan) to construct the optical tweezers. The microscope stage was replaced by a piezoelectric stage controlled by a stage controller (MDR14-CA-2.5; SIGMAKOKI, Tokyo, Japan) and a joystick (JS-300; SIGMAKOKI). The irradiated laser beam was concentrated as a finite optical system using a plano-convex lens supported by an optical cage system (SIGMAKOKI). The concentrated laser beam was inserted into the microscope and focused by an objective lens (CFI Apochromat TIRF 100XC Oil; Nikon, Tokyo, Japan). The actual laser power was measured using a power meter (FieldMaxII; COHERENT, Santa Clara, CA) without the objective lens.

### Force measurements

The cell suspension was mixed with 0.5 mM Sulfo-NHS-LC-LC-biotin (Thermo Fisher Scientific, Waltham, MA) as the final concentration and incubated for 15 min at room temperature (RT). The cell suspension was centrifuged at 12,000 × *g* for 10 min, washed with 10 mM HEPES buffer (pH 7.4) containing 150 mM NaCl, centrifuged again, washed with HEPES buffer containing 10% non-heat-inactivated horse serum (Gibco; Thermo Fisher Scientific) and 20 mM glucose, passed through a 0.45-μm pore size filter and incubated for 15 min at RT. The cell suspension was inserted into a tunnel chamber, which was assembled by taping coverslips cleaned with saturated ethanolic KOH and precoated with 100% non-heat-inactivated horse serum for 60 min and 10 mg/ml bovine serum albumin (Sigma-Aldrich, St. Louis, MO) in HEPES buffer for 60 min at RT. The tunnel chamber was washed with HEPES buffer containing 20 mM glucose and incubated at 37°C on optical tweezers equipped with a thermo plate (MATS-OTOR-MV; Tokai Hit, Shizuoka, Japan) and a lens heater (MATS-LH; Tokai Hit). Avidin-conjugated beads in the HEPES buffer containing 20 mM glucose and 0.2–0.3 mM of 3’-*N*-acetylneuraminyllactose (SL) were sonicated and inserted into the tunnel chamber. The avidin-conjugated beads were prepared as previously described (35). The bead movements were recorded using a charge-coupled device (CCD) camera (LRH2500XE-1; DigiMo, Tokyo, Japan) at 30 frames per second and analyzed by displacement of up to 200 nm from the trap center (the linear range of the laser trap) using ImageJ 1.43u (http://rsb.info.nih.gov/ij/) and IGOR Pro 6.33 J and 8.02 J (WaveMetrics, Portland, OR) (34, 35, 62).

### Binding and gliding analyses

Cultured cells were washed with buffer in a culture flask, and then washed with HEPES buffer containing 10% non-heat-inactivated horse serum (Gibco; Thermo Fisher Scientific) and 20 mM glucose. Cultured cells were scraped off the culture flask, passed through a 0.45-μm pore size filter, and incubated for 15 min at RT. The cell suspension was then inserted into the tunnel chamber. The tunnel chamber was washed with HEPES buffer containing 20 mM glucose, incubated at 37°C on an inverted microscope (IX83; Olympus) equipped with a thermo plate and lens heater, observed by phase-contrast microscopy at 37°C, and recorded with a CCD camera (DMK 33UX174; The Imaging Source Asia Co. Ltd., Taiepi City, Taiwan). Then, the tunnel chamber was washed with HEPES buffer containing 20 mM glucose and various concentrations of SL. Video data were analyzed using ImageJ 1.43u and IGOR Pro 6.33 J.

### Genome sequencing and variant analyses

Frozen stocks of cells were plated on Aluotto medium and isolated as previously described (63). Genomic DNA was isolated and sequenced using MiSeq (Illumina, San Diego, CA), as previously described (34). Sequence read mapping and variant detection were performed using the CLC Genomics Workbench (QIAGEN, Hilden, Germany).

## Acknowledgments

We thank Dr. Tsuyoshi Kenri at the National Institute of Infectious Diseases and Dr. Ikuko Fujiwara at Osaka City University for helpful discussions, and Dr. Shigeyuki Kakizawa at the National Institute of Advanced Industrial Science and Technology (AIST) for supporting the genome sequence analyses. This work was supported by Grants-in-Aid for Scientific Research: (A) (MEXT KAKENHI Grant Numbers JP24390107, JP17H01544) and by JST CREST (Grant Number JPMJCR19S5, Japan). M. Mizutani is the recipient of a Research Fellowship of the Japan Society for the Promotion of Science (18J15362).

## Notes

### Competing Interest Statement

The authors have declared no competing interest.

### Summary of Updates

New data has been added. Supplemental files updated.

