## Supplementary material for "Force and stepwise movements of gliding motility in human pathogenic bacterium *Mycoplasma pneumoniae*": Fig S1-3, Table S1,

Running title: Force and step of *Mycoplasma pneumoniae* gliding

Keywords: Motility, Optical tweezers, class *Mollicutes*, Infection

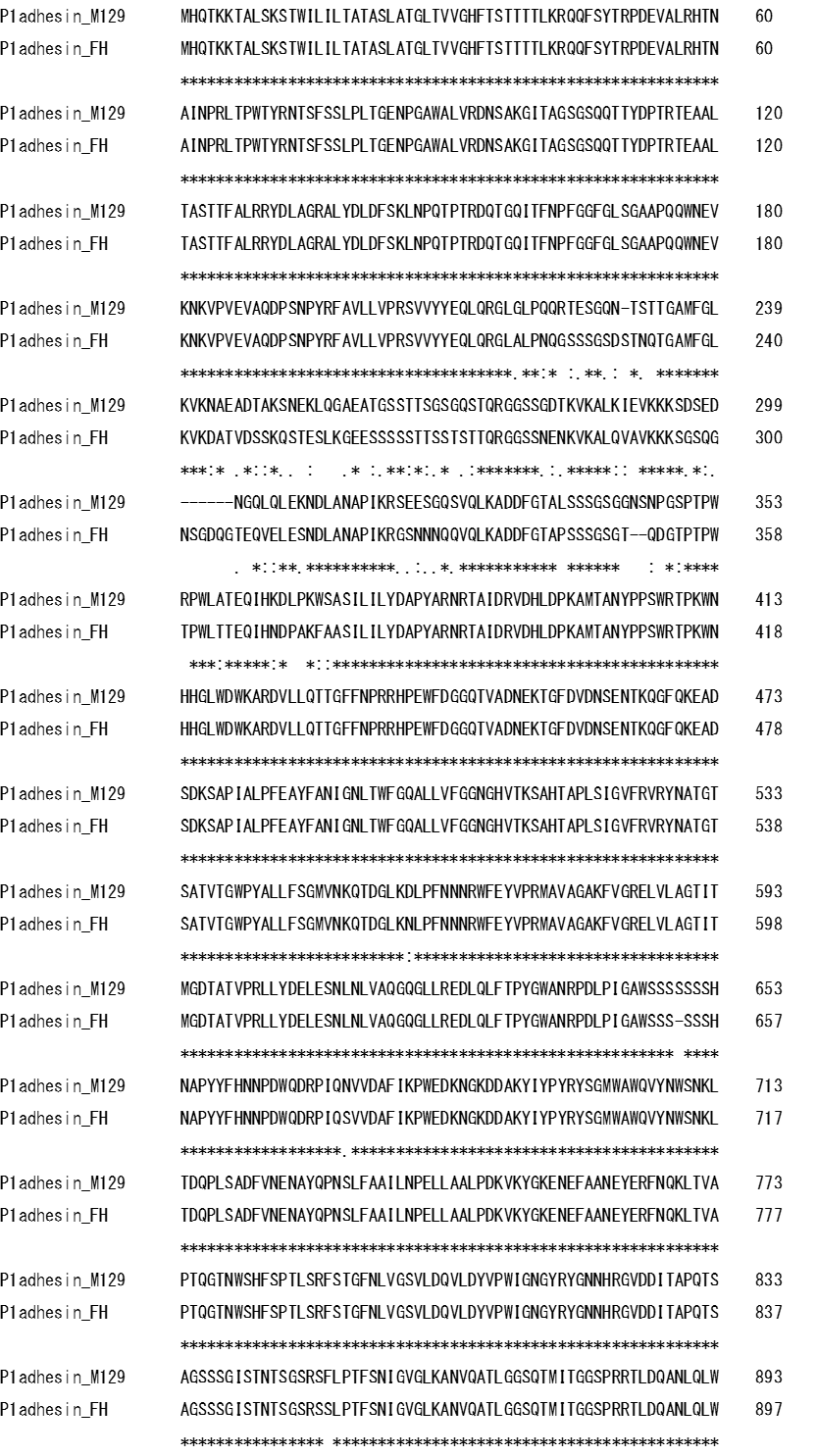

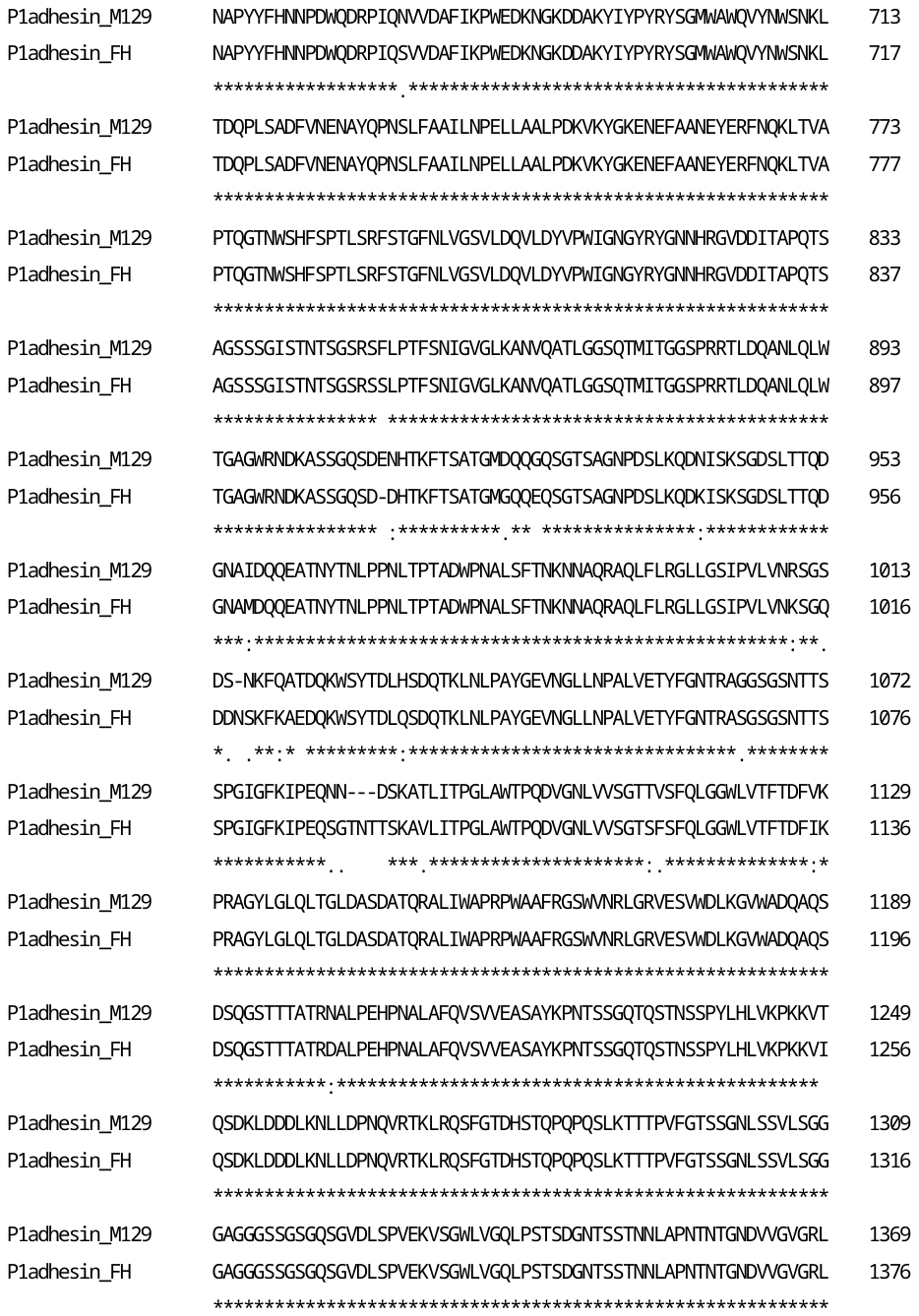

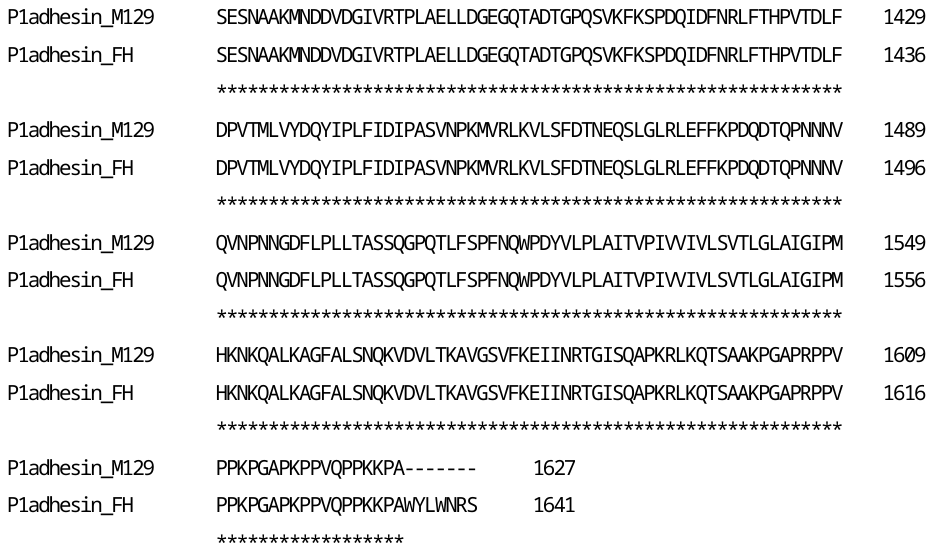

**Fig. S1. Multiple sequence alignments for P1 adhesin of M129 and FH strains.** The symbols “*” “:” “.” indicate fully conserved residue, conservation between groups of strongly similar properties, and conservation between groups of weakly similar properties, respectively.

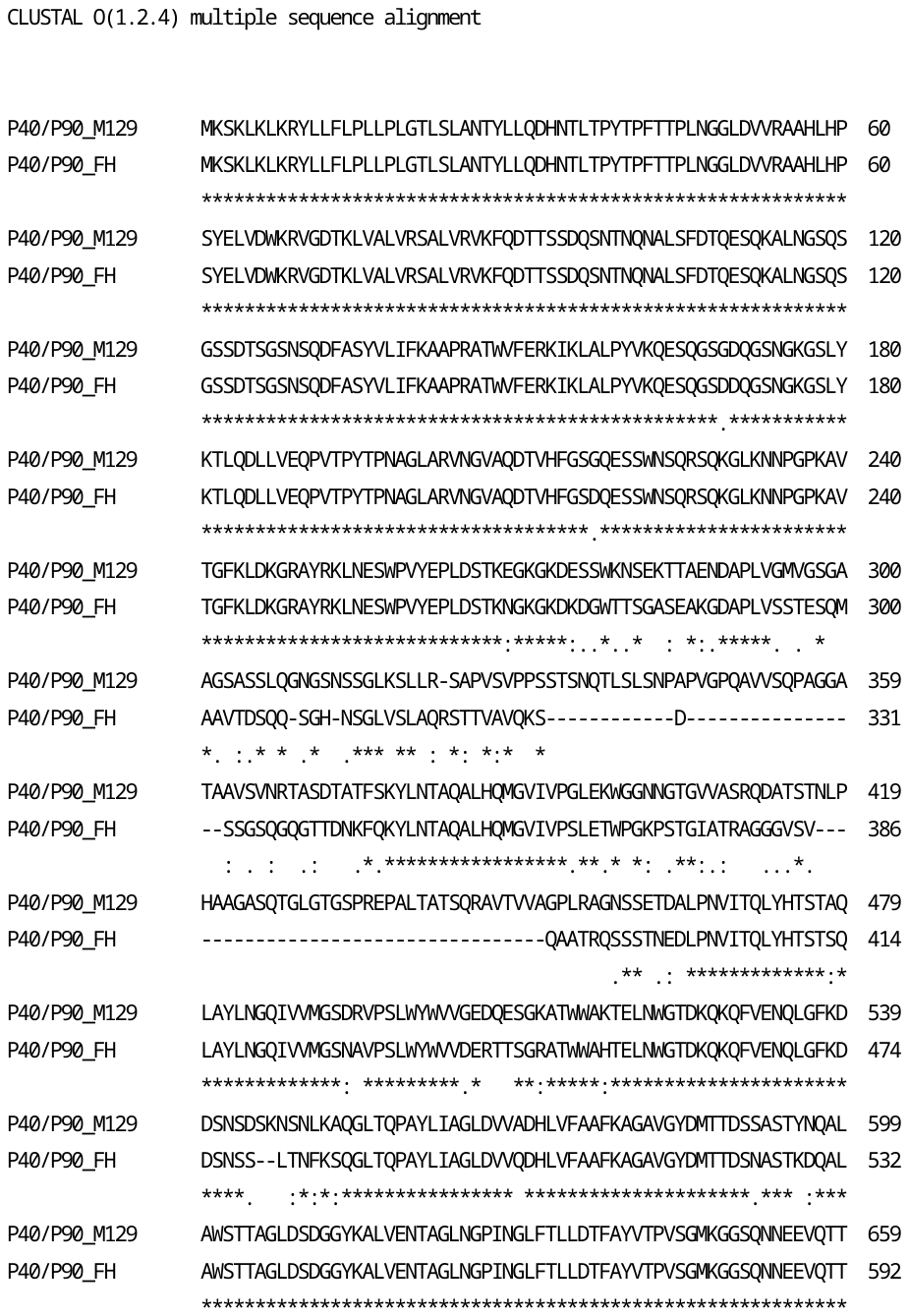

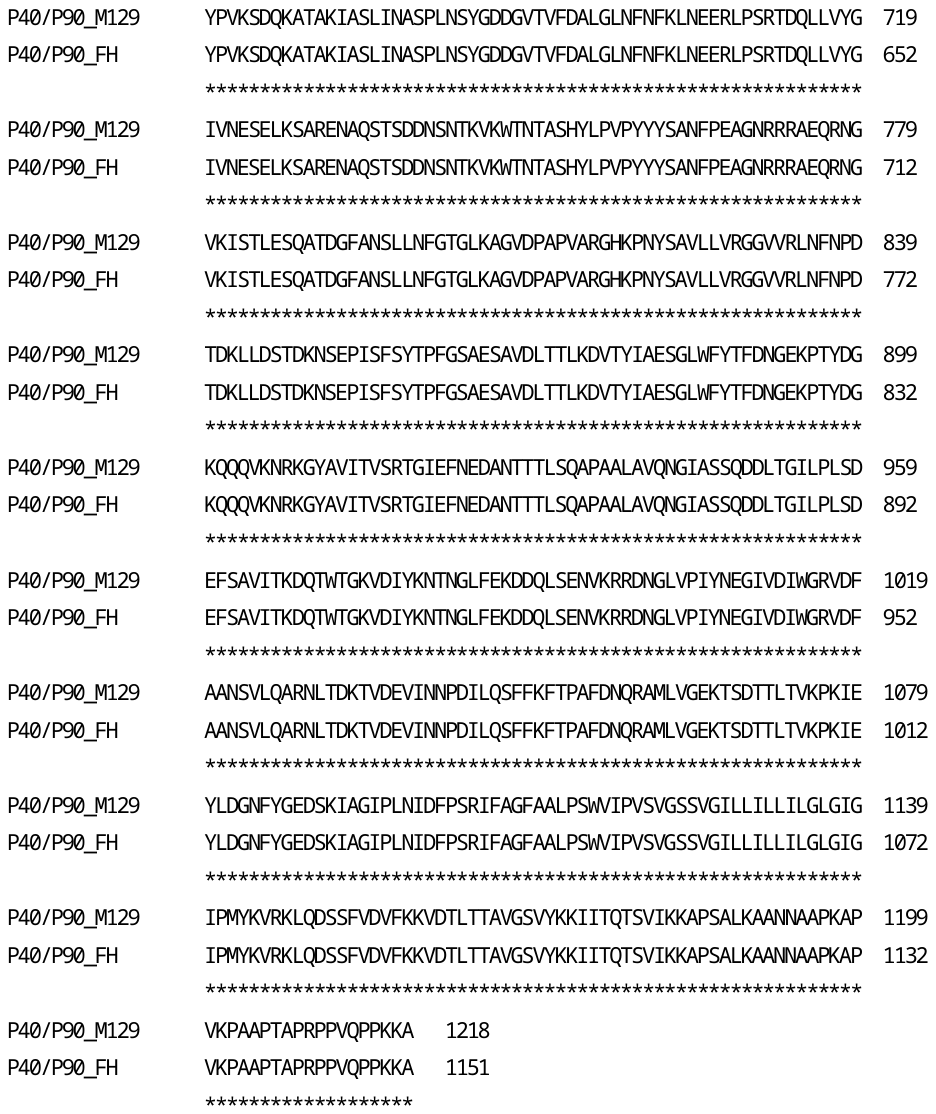

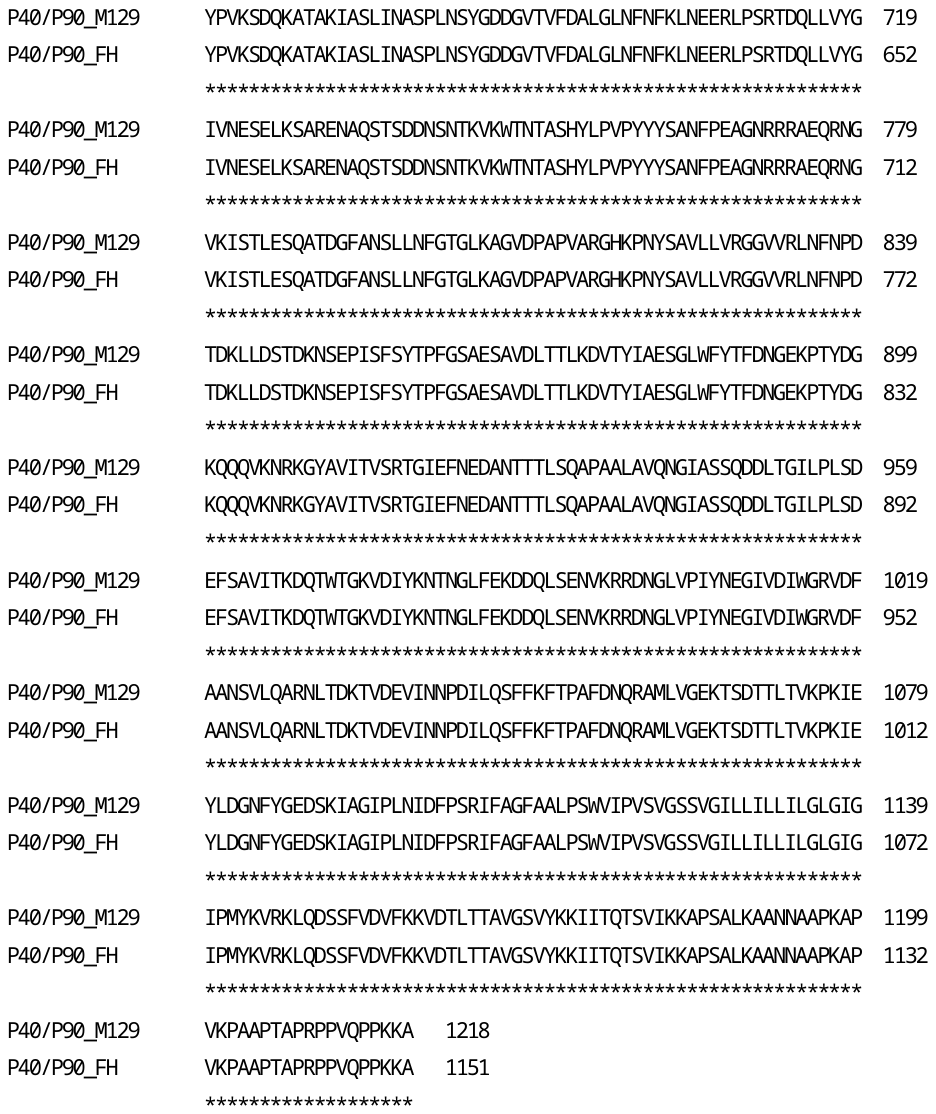

**Fig. S2. Multiple sequence alignments for P40/P90 of M129 and FH strains.** The symbols “*” “:” “.” indicate fully conserved residue, conservation between groups of strongly similar properties, and conservation between groups of weakly similar properties, respectively.

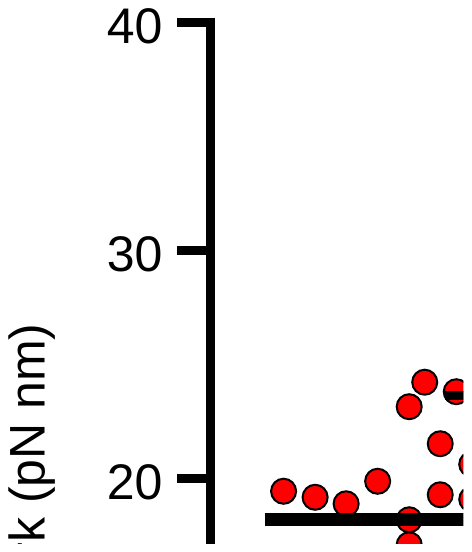

**Fig. S3. Work performed by stepwise movements.** The scatter dot plot of works calculated from individual steps is shown with average (thick line) and standard deviation (thin lines).

**Table S1. Amino acid variations of gliding related proteins in M129 and FH strains.**

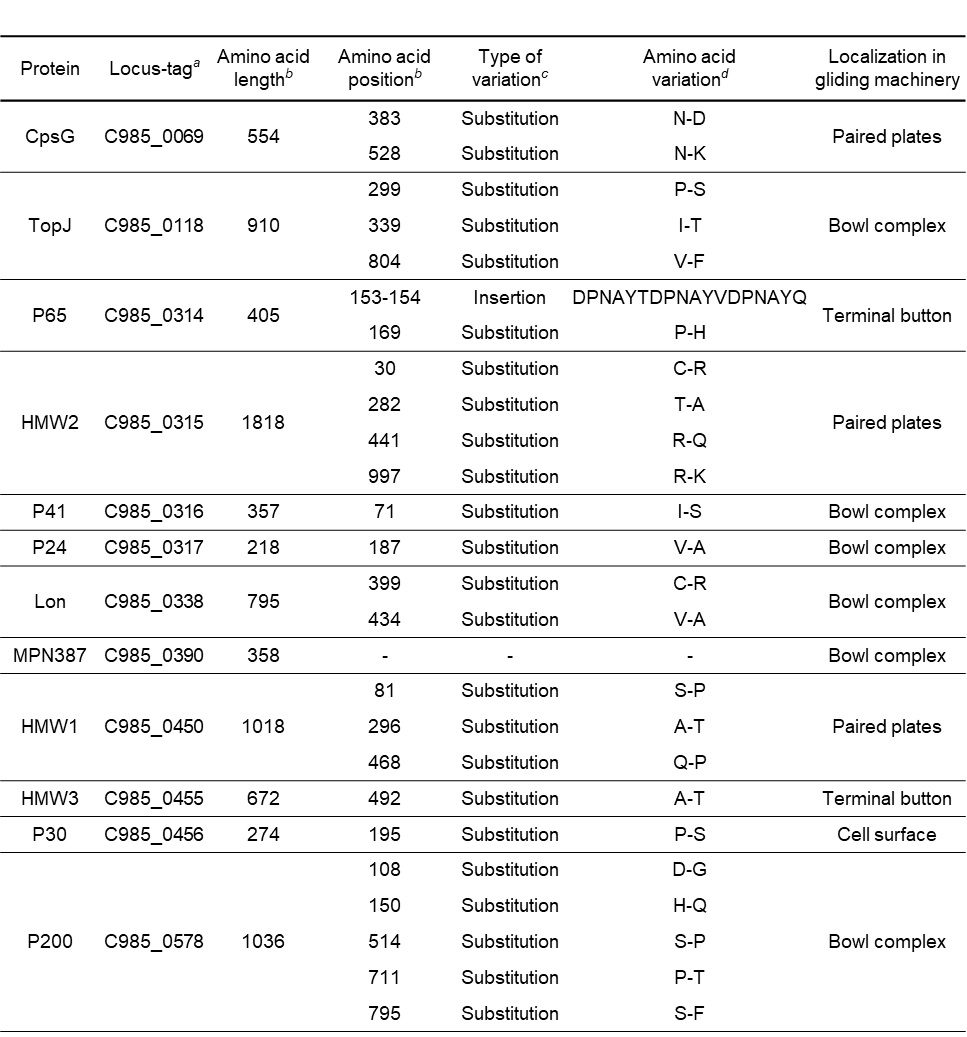

*^a^* Locus-tag is as in the sequence under accession number CP003913 for M129-B7.

*^b^* The values are for M129.

*^c^* The variations were detected in FH against M129.

*^d^* The amino acids are written the order M129-FH.

**Movie S1 (separate file). Stall force measurement in M129 strain.** A polystyrene bead was attached to the back end of cell body. The cell pulled the bead from trap center of optical tweezers. The video was played at 5 × speed.

**Movie S2 (separate file). Stall force measurement in FH strain.** A polystyrene bead was attached to the back end of cell body. The cell pulled the bead from trap center of optical tweezers. The video was played at 5 × speed.

**Movie S3 (separate file). Gliding movement of M129 strain cells.** Cells bound to the SOs-coated glass surface were observed by phase-contrast microscopy. The video was played at 5 × speed.

**Movie S4 (separate file). Gliding movement of FH strain cells.** Cells bound to the SOs-coated glass surface were observed by phase-contrast microscopy. The video was played at 5 × speed.

**Dataset S1 (separate file).** Variant analyses of whole genome for our M129 and FH strains against the reported M129-B7 and FH strain.
